# Characterisation of *Skoliomonas* gen. nov., a haloalkaliphilic anaerobe related to barthelonids (Metamonada)

**DOI:** 10.1101/2024.03.12.584707

**Authors:** Yana Eglit, Shelby K. Williams, Andrew J. Roger, Alastair G.B. Simpson

**Affiliations:** Department of Biology, and Institute for Comparative Genomics, Dalhousie University, Halifax, NS, Canada; Department of Biochemistry, and Institute for Comparative Genomics, Dalhousie University, Halifax, NS, Canada

**Keywords:** protists, anaerobes, protist diversity, phylogenetics, alkaliphiles

## Abstract

Metamonads are a large and exclusively anaerobic clade of protists. Additionally, metamonads are one of the three clades with a proposed ‘excavate’ ancestral cell morphology, characterised by a conspicuous ventral groove often accompanied by a posterior flagellum with a vane. Here, we characterise four isolates of an anaerobic bacterivorous flagellate from hypersaline and alkaline soda lake environments, which represents a novel clade. Small subunit ribosomal RNA (SSU rRNA) gene phylogenies support recent phylogenomic analyses in placing this clade as the sister group to *Barthelona* spp., a lineage that is itself sister to or deeply branching within Fornicata (Metamonada). The cells have a distinctive morphology comprised of a hunchbacked cell body with a narrow twisting ventral groove ending in a large opening to a conspicuous cytopharynx curving up the dorsal side of the cell. The right margin of the groove is defined by a thin ‘lip’ that twists slightly to the left towards the posterior. The posterior of the cell ends in a spike up to half a cell body long. The posterior flagellum bears a wide ventral-facing vane. One isolate forms cysts with a complex wall and a single plug. The narrow ventral groove and elongate cytopharynx are shared with barthelonids. We describe one isolate as *Skoliomonas litria*, gen. et sp. nov. Further investigation of mitochondrial-related organelles (MRO) in *Skoliomonas* spp. and detailed ultrastructural studies would be important to understanding the evolution of adaptation to anaerobic conditions in Metamonads—especially fornicates—as well as the evolution of the ‘excavate’ groove.

## Introduction

Adaptation to life at low-to-no oxygen is widely distributed across the tree of eukaryotes (Gawryluk and Stairs, 2021; Leger et al., 2019). Yet, many described species belong to just one clade, the metamonads (Metamonada), a diverse group of protists that consists entirely of anaerobes (Burki et al., 2020). All studied metamonads possess substantially modified mitochondria, referred to as mitochondrion-related organelles (MROs) (Leger et al., 2019), or lack mitochondrial organelles altogether (Karnkowska et al., 2016). Some lineages possess further unusual biological features like missing key components of the canonical division machinery (Salas-Leiva et al., 2021) or are surprisingly symbiotically complex amoebae (Jerlström-Hultqvist et al., 2023; Stairs et al., 2021; Táborský et al., 2017).

The best known metamonad taxa—Parabasalia, Diplomonadida and Oxymonadida—are each ancestrally commensals, symbionts or parasites of animals (Archibald et al., 2017; Céza et al., 2022; Mazancová et al., 2023; Novák et al., 2023). Parabasalids and Diplomonadids in particular include medically- and economically-significant parasites like *Trichomonas* or *Giardia* species (respectively), which are extensively studied. Yet, most of the phylogenetic diversity of Metamonada is free-living and not as well-investigated. Most free-living metamonads have two or four flagella and possess an ‘excavate’-type feeding groove. This groove is associated with the posterior flagellum, which beats to generate a feeding current (Simpson, 2003; Simpson and Patterson, 1999; Suzuki-Tellier et al., 2024). This flagellum typically has a vane; the beating of this flagellum draws prey bacteria into the groove and toward the distal end, where phagocytosis occurs, while computational modelling indicates that the combination of vane and groove increases the magnitude of the feeding current (Suzuki et al., 2023; Suzuki-Tellier et al., 2024).

The majority of described free-living metamonad lineages are so-called ‘*Carpediemonas*-like organisms’ (CLOs). These were characterised relatively recently, and are related to diplomonads, with the clade containing CLOs and diplomonads (and the more obscure retortamonads) known as Fornicata (Park et al., 2010, 2009a; Simpson et al., 2002; Yubuki et al., 2016, 2013). However, significant novel free-living lineages are still being found; for example, *Anaeramoeba*, a genus of amoebae found in anaerobic environments (Táborský et al., 2017), was determined to be a deep-branching free-living relative of parabasalids (Stairs et al., 2021). Recently, the first molecular data were obtained for the obscure anaerobic flagellate *Barthelona* (Bernard et al., 2000), and these ‘barthelonids’ were determined to be sister to fornicates (Yazaki et al., 2020). Understanding the diversity of free-living metamonads is important for tracing the evolutionary history of this clade, including the origins of parasitism and innovations shaping anaerobic metabolism and MRO evolution.

Soda lakes, a subset of alkaline lakes rich in carbonate ions, are a unique and often productive habitat with a distinctive prokaryote and large-eukaryote biodiversity (Pirlot et al., 2005; Yasindi and Taylor, 2006; Zorz et al., 2019). They are typically found as endorrheic water bodies in regions with little sedimentary rock exposure (Jones et al., 1998; Schagerl and Renaut, 2016). Examples include lakes in the East African Rift Valley (Schagerl and Burian, 2016) and in interior mountain ranges on the west coast of North America (Rojas et al., 2018; Sorokin et al., 2007; Zorz et al., 2019). Many of these lakes fluctuate extensively in water level and salinity throughout the seasons (Schagerl and Burian, 2016). Thus, not only do their inhabitants face high pH accompanied by unusual chemical environments, but also variable salinity that may range from nearly freshwater to nearly saturated. Additionally, some of the shallower lakes can reach high temperatures (Jones et al., 1998; Schagerl and Burian, 2016). Thus, many microbes thriving in these environments are polyextremophiles (Bowers et al., 2009; Oren, 2013).

Studies of microbial diversity in soda lakes have largely focused on prokaryotes or, in some cases, eukaryotic algae (e.g. Rojas et al., 2018; Sorokin et al., 2015, 2014; Sousa et al., 2015; Zorz et al., 2019). With the exception of some extensive work on ciliates and their connection to bioproductivity in East African Rift Valley lakes (Yasindi and Taylor, 2006, 2016, 2020; Schagerl and Burian, 2016), little attention has been given to heterotrophic protists in these lakes. Two recently described species, *Kaonashia insperata* (Weston et al., 2023) and *Neocolponema saponarium* (Gigeroff et al., 2023), are eukaryotrophic aerobes. To our knowledge, eukaryotic haloalkaliphilic anaerobes have not been documented in soda lakes to date.

Here, we obtained four isolates of a previously unidentified group from three geographically distinct soda lakes. The organisms resemble free-living metamonads in being anaerobic flagellates with a ventral feeding groove associated with the posterior flagellum. Ribosomal RNA (rRNA) gene phylogenies confirm that they represent a clade that does not branch within any of the previously identified major categories of metamonads. We describe one isolate (TZLM1-RC) as the type of a new genus, *Skoliomonas litria* n. gen. n. sp. and refer to the group collectively as ‘skoliomonads’.

## Methods

### Isolation and cultivation

The metamonad strains were isolated from samples obtained from alkaline (pH ∼10) hypersaline (up to 160 ppt salinity) sediments from Lake Manyara, Tanzania (TZLM1 and TZLM3; September, 2016; 3°37’01.7”S, 35°44’22.8”E), Goodenough Lake, BC, Canada (GEM; May, 2017, 51° 21’ 54” N, 121° 47’ 41” W) and Soap Lake, WA, US (Soap20A; December, 2020, 47°23’28.2”N, 119°29’11.9”W). An additional isolate from the latter locality, Soap18-RC., was cultivated in 2018 from a 30 ppt sample. Soap18-RC was indistinguishable from Soap20-RC in SSU rRNA gene sequence (see below). Isolates were enriched in 15mL conical tubes filled to 12mL with 160ppt (TZLM1), 50ppt (GEM), 20ppt (Soap20), and 5ppt (TZLM3) versions of medium “CR” (Gigeroff et al., 2023) (KH_2_PO_4_ 0.2, MgCl_2_ 0.1, KCl 0.2, NH_4_Cl 0.5, NaCl 100, Na_2_ CO_3_ 68, NaHCO_3_ 38 g•L^-1^; adapted from (Mesbah et al., 2007), roughly matching the source salinity level, with sterile barley grains and/or 1-3% (final concentration, v/v) LB added to support bacterial growth and maintenance of low oxygen conditions. Cultured isolates were purified by dilution over repeated transfers. Soap20A-RC was passaged through 2 rounds of 100 ppt “CR” medium before reducing salinity to 50 ppt. Purified TZLM1-RC, Soap20A-RC, GEM-RC and TLZM3-RCL lines were maintained in medium “CR” diluted to 100, 50, 50, and 5 ppt, respectively, with four sterile wheat grains /12 mL (and an additional 1% and 3% v/v LB for GEM-RC and TZLM1-RC, respectively) to sustain bacterial growth, transferred every 2-3 months except for TZLM3-RCL, which was transferred every other week. Alternative media supporting relatively high growth were established for three isolates, as follows: for TZLM1-RC: CR40 + 3% v/v LB; for GEM-RC: CR40 + 1% v/v LB + 1 sterile wheat grain / 3 mL; for TZLM3-RCL: CR25 + 1% v/v LB + 1 sterile wheat grain / 3 mL. Resazurin (Karakashev et al., 2003) was used early in the enrichment and cultivation process to confirm anoxic conditions.

#### Microscopy

Unless stated otherwise, imaging was done with a Zeiss Axiovert 200M microscope with DIC optics, a 100x objective, and a Zeiss Axiocam HRc camera (Carl Zeiss AG, Oberkochen, Germany) on live cells maintained in a vaseline-sealed coverslip-slide ‘chamber’ for around 24h. To measure cell sizes, we observed early stationary phase cultures 2 weeks (TZLM1-RC, GEM-RC, TZLM3-RCL) or 6 weeks (Soap20A-RC) old, grown at 21°C under culturing conditions described above (RC: CR40 + 3% v/v LB, GEM-RC: CR40 + 1% v/v LB + 1 sterile wheat grain / 3 ml, SP20A-RC: CR50 + 1 sterile wheat grain / 3 ml, RCL: CR25 + 1% v/v LB + 1 sterile wheat grain / 3 ml). Fluid was taken from the bottom of the tube/flask without agitation, placed on glass slides, capped with a coverslip (without vaseline and with no delay before imaging), and cells exhibiting flagellar beating were imaged under phase contrast using a 40x objective and 1.6x optovar. Sizes were measured from captured digital images using FIJI (Rasband, 1997; Schneider et al., 2012). Videos, from which some still images were taken, were recorded on a Nikon Eclipse Ti2 inverted microscope and Digital Sight 10 camera (Nikon) with DIC optics and a 100x objective with or without a 1.5x tube lens.

Preliminary transmission electron microscopy (TEM) images corroborating light microscopy results were obtained by pelleting cultures of TZLM1-RC (in CR160) and TZLM3-RCL (in CR5) via centrifugation at 13k×*g* and subjecting them to fixation in 5% glutaraldehyde in CR50 medium or 2.5% glutaraldehyde in deionised water, respectively, for an hour, followed by a 30 min postfix in 1% OsO_4_ in CR25 medium for TZLM1-RC and in deionised water for TZLM3-RCL. Cells were dehydrated in an acetone series (33%, 50%, 75%, 90%, 100% v/v in water) and embedded in SPI-PON 812 resin. A subsample of this resin was mounted on slides to make permanent mounts of type material. The resin blocks were then sectioned on a Leica EM UC7 ultramicrotome and unstained sections imaged with the Gatan 832 SC1000 camera on FEI Tecnai-12 electron microscope.

#### DNA extraction, amplification and sequencing

Genomic DNA for each of the cultured isolates was extracted using the Qiagen DNeasy Blood & Tissue kit (CAT#69504), except for GEM-RC, which was obtained using a combination of phenol:chloroform and CTAB extraction protocols, the QIAGEN Genomic-tip kit, and the QIAGEN MagAttract HMW DNA kit, further detailed in (Williams et al., 2023). Partial SSU rDNA sequences for TZLM1-RC and Soap18-RC, and GEM-RC and Soap20A-RC were amplified using EukA (5l⍰-AACCTGGTTGATCCTGCCAGT-3l⍰) and EukB (5’-TGATCCTTCTGCAGGTTCACCTAC-3’) primers (Medlin et al., 1988) at 60°C and 58°C annealing temperatures, respectively. The partial SSU rDNA sequence for TZLM3-RCL was amplified using 82F (5l⍰-GAAACTGCGAATGGCTC-3l⍰) and 1498R (5’-CACCTACGGAAACCTTGTTA-3’) at 55°C annealing temperature. All PCR reactions were run with 35 cycles under the following program: 98°C for 2 min; 98°C for 30 s, annealing temperature for 1 min, and 72°C for 2 min 30s; with a final elongation step at 72°C for 10 min. The amplified product was Sanger sequenced at Génome Québec, with the reads trimmed and assembled in Geneious Prime 2022.1.1 (https://www.geneious.com). For phylogenetic analyses (see below), the partial and lower quality PCR-derived TZLM3-RCL SSU rDNA sequence was replaced by one extracted from the TZLM3-RCL genome assembly (Williams et al., 2023) using barrnap (Seeman, 2018).

#### Ribosomal gene phylogenies

Ribosomal gene sequences were aligned to a manually curated eukaryote-wide alignment in SeaView (Gouy et al., 2010) using profile alignment via MUSCLE (Edgar, 2004), trimmed with a gblocks (Castresana, 2000) masking further curated by hand, with 1321 sites retained and 219 taxa. SSU and LSU rRNA gene sequences corresponding to the same operon were extracted from genomic data of skoliomonad isolates TZLM1-RC, GEM-RC, and TZLM3-RCL as well as barthelonid PCE from (Williams et al., 2023), then added to a concatenated SSU-LSU rDNA alignment derived from (Eglit et al., 2024), which was then trimmed with gblocks and manually edited to a total of 3064 sites and 141 taxa. For each dataset, a maximum likelihood phylogeny was inferred in RAxML v8.2.6 (Stamatakis, 2014) under the GTR+Г model with 50 starting trees and 500 non-parametric bootstraps.

An uncorrected distance matrix was calculated for the SSU rRNA gene sequences used in the global SSU rDNA phylogeny using SplitsTree4 (Huson and Bryant, 2006) and visualised as a heatmap using the Seaborn package (Waskom, 2021). Final tree figures were rendered using the Ete3 toolkit (Huerta-Cepas et al., 2016) and finished in Inkscape v1.2 (The Inkscape Project, 2022).

## Results

### Light Microscopy

Skoliomonad cells have a rounded anterior end and a pointed posterior, with a dorsal hump and a flattened ventral side containing a major groove (Fig 1 A-K, O). The posterior ends in a spike that can be up to half the length of the cell body proper (Fig 1 A, D, F, H). The left side of the cell contains the bulk of the cytoplasm, including the nucleus and numerous large digestive vacuoles. Long prey bacteria can be found inside vacuoles along the dorsal side of the cell (Fig 1 C, E, H). The right side is comprised almost entirely of the right edge of the groove (see below). At least one isolate, TZLM3-RCL, forms cysts (Fig 1 L,M; Fig 2 T).

**Figure 1.**
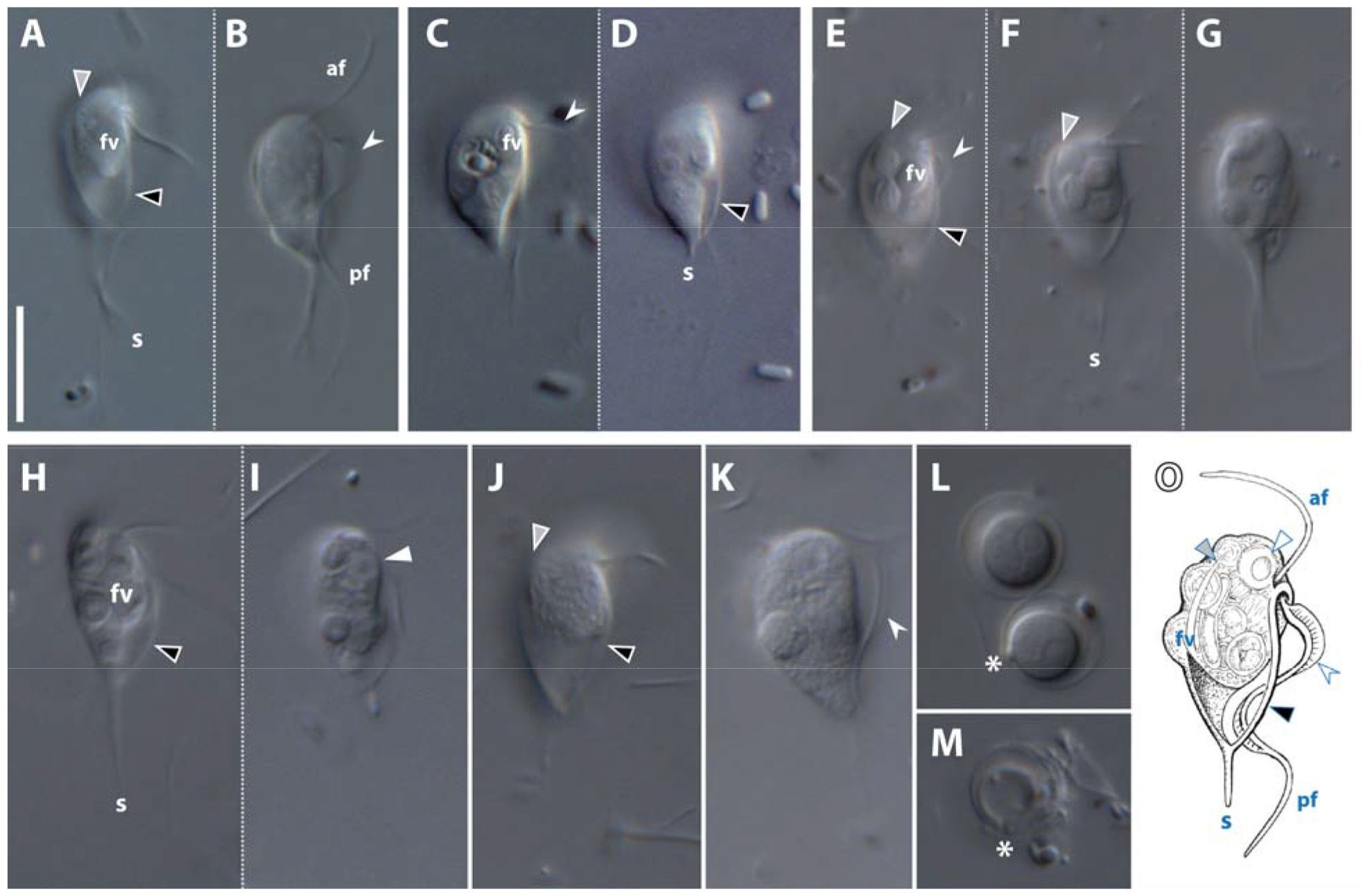
Differential interference contrast (DIC) images showing the general morphology of *Skoliomonas litria* n. gen., n. sp. TZLM1-RC (A-B), *Skoliomonas* sp. Soap20-RC (C-D), *Skoliomonas* sp. GEM-RC (E-F), and *Skoliomonas* sp. TZLM3-RCL (H-M). A) Optical section showing the right edge of the cell showing the lip (black arrowhead) and the cytopharynx (grey arrowhead). B) Section of the same cell further to its left. The vane on the posterior flagellum is indicated by a barbed arrowhead. C-D) Optical sections of the same cell showing general morphology of Soap20A-RC. E-G) Optical sections of the same cell of isolate GEM-RC showing the general morphology, including a section deeper into the cell’s left showing a portion of the ventral groove opening (G). H-I) Optical sections of the same cell of TZLM3-RCL showing the general morphology of a skinnier cell. J-K) Two different cells of isolate TZLM3-RCL showing bigger individuals with the same key morphological features. L-M) Complete (L) and excysted (M) cysts of TZLM3-RCL. Asterisk indicates the cyst plug location. Note the inner and outer wall and the space between them. O) General diagram of a skoliomonad cell. af, anterior flagellum; pf, posterior flagellum; fv, food vacuole; s, (posterior) spike; grey arrowhead, cytopharynx; black arrowhead, ‘lip’ of the right sheet; barbed arrow, flagellar vane; white arrow, nucleus; asterisk, cyst plug. Scale bar is 10 μm for all micrographs.

**Figure 2.**
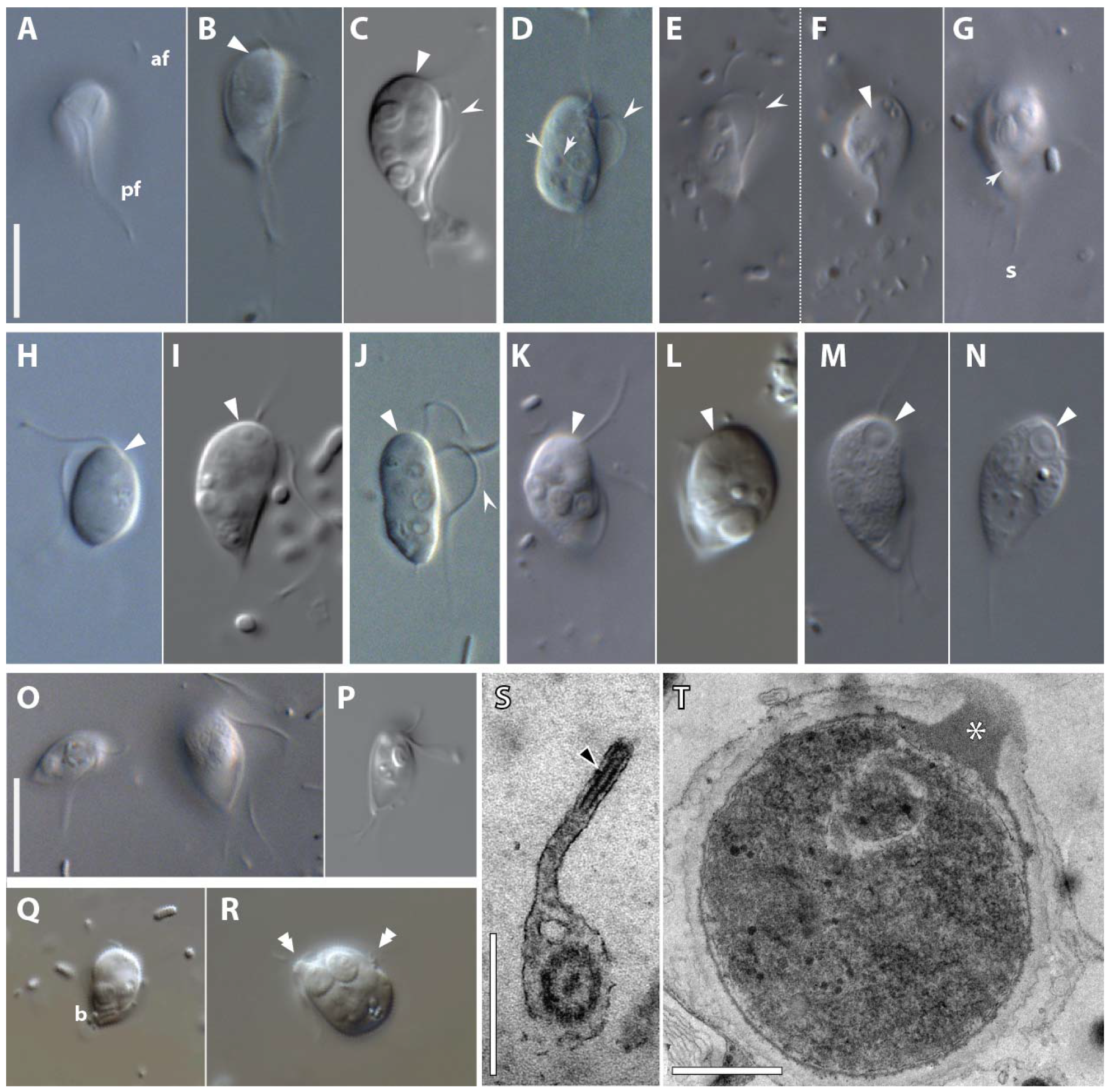
Additional light and transmission electron microscopy images for the four *Skoliomonas* sp. isolates. A-G) Further cell morphology details in TZLM1-RC (A-C), Soap20-RC (D), and GEM-RC (E-G). A) Optical section of the ventral groove and posterior flagellum. B) Optical section further towards to the left side of a different cell showing the anterior extent of the recurrent cytopharynx. C) Optical section through the leftmost part of another cell. D) Flattened cell showing the flagellar vane (barbed arrowhead) as well as the robust walls of the cytopharynx (arrows). E) View of the posterior flagellum vane. F) Oblique optical section through the anterior-dorsal side of the cell. G) Grazing section of the right side of the cytopharynx. H-N) nuclei in the four isolates: TZLM1-RC (H-I); Soap20-RC (J) GEM-RC (K-L); TZLM3-RCL (M,N). Note subtle differences in morphology. O-P) Additional images of TZLM3-RCL, showing size variation in a single frame (O) and detail of the smaller morph (P). Q-R) Images of GEM-RC showing early stages of bacterial ingestion (Q), and a dividing cell with two sets of flagella (double arrowheads)(R), both stills from Supplementary video (XXX). TEM section through the posterior flagellar vane in TZLM1-RC. T) TEM section through a cyst of TZLM3-RCL showing the complex walls and the plug (asterisk). af -- anterior flagellum; pf -- posterior flagellum; white arrowhead -- nucleus; barbed arrowhead -- flagellar vane; arrows -- cytopharynx walls; s -- (posterior) spike; b -- bacterium; double arrowheads -- duplicated flagellar sets; black arrowhead -- intraflagellar structure supporting the vane; asterisk -- plug. Scale bars: 10 μm in A and O; 500 nm in S; 1 μm in T. A-R are at the same scale.

Two flagella of different lengths insert subapically facing the ventral side of the cell at a slightly obtuse angle (Fig 1 A, B). A structure extends from there towards the posterior of the cell and forms the left side of a subtle ventral groove. A large conspicuous ‘lip’ extends from the flagellar insertion along the right side of the groove, to the base of the posterior spike (Fig 1 A, H).

The anterior flagellum is approximately a cell-length long and points forward (Fig 1 B; Fig 2 K). The posterior flagellum is two cell-lengths long and features a spectacularly broad vane about 1 μm wide along the length of the groove directed ventrally—that is, away from the cell body (Fig 1B, E, K; Fig 2 C, E, J). The existence of a single vane is supported by preliminary TEM, which also shows an internal structure supporting the distal edge of the vane (Fig 2 S). A prominent structural element initiates slightly to the left of the posterior flagellum, wraps around anterior side of it and forms a lip delineating the right edge of the groove (Fig 1A, E, H). The groove curves gently to the right as it extends down the cell, generally resembling the opening of an olive snail (Fig 1G). The distal end of the groove features a large opening to a substantial scythe-shaped cytopharynx immediately underneath the right lip, with the breadth of the opening comprising nearly half of the length of the groove (Fig 1A, E, F, J; 2B). The cytopharynx narrows as it extends along the dorsal side back towards the apex of the cell. It is supported by a robust intracellular structure, as seen in cells tightly flattened by a coverslip (Fig 2D).

The nucleus (Fig 2 H-N) is anteriorly situated, to the right of the flagellar insertion, and has a conspicuous eccentric nucleolus in TZLM1-RCL (Fig 2M-N). The nucleus is more subtle in the other isolates (Fig 2 H-L). There are numerous bacteria in vacuoles, presumably prey (Fig 1A, C, E). The food vacuoles may distort the shape of the cell, particularly on its left and dorsal sides (Fig 1B; Fig 2K). Some food vacuoles are elongate and contain intact long bacteria, but most are rounded— presumably representing later stages of digestion. Ingestion has been observed at the opening to the cytopharynx, with prey bacteria taken up along the cytopharynx on the dorsal side of the cell (Fig 2Q).

The cells move using both flagella in a vibrating motion, with the posterior directed backwards in a large amplitude wave as the cell gently rotates, and the anterior flagellum pointing slightly forward. On a slide, the cells spend significant time swimming along surfaces in circular paths. Cells were not found to attach and filter currents in chamber slide setups; however, in immediately-imaged conventional preparations individuals of TZLM3-RCL were found to stick to the surface with the tip of the spike, later capable of detachment. In similar preparations, cells of TZLM1-RC appeared to attach with the cell surface, presumably an artefact. This was not observed in either TZLM3-RCL nor GEM-RC under similar conditions.

Cells are 8.1 – 13.6μm (mean: 10.6±1.5μm), 7 – 13.5μm (mean 10.1±1.3μm), 9 – 15μm (mean 11.2±1.2μm), and 8.9 – 12.7 μm (mean: 10.5±1.0μm) long, excluding the tail spike, and 4.2 – 6.8 μm (mean: 5.8±0.7 μm), 4.5 – 8 μm (mean: 5.7±0.7μm), 4.6 – 9.5 μm (mean: 5.9±1.1μm), and 4.2 – 6.8 μm (mean: 5.3±0.7μm) wide for TZLM1-RC, TZLM3-RCL, GEM-RC, and Soap20-RC, respectively (see Table 1).

**Table 1.** Cell size metrics for flagellates of *Skoliomonas* spp. isolates.

|  | Mean<br>length<br>(µm) | stdev<br>length<br>(µm) | mean<br>width<br>(µm) | stdev<br>width<br>(µm) | min length<br>(µm) | max length<br>(µm) | min width<br>(µm) | max width<br>(µm) | n |
| --- | --- | --- | --- | --- | --- | --- | --- | --- | --- |
| **TZLM1-RC** | 10.6 | 1.5 | 5.8 | 0.7 | 8.1 | 13.6 | 4.2 | 6.8 | 33 |
| **TZLM3-RCL** | 10.1 | 1.3 | 5.7 | 0.7 | 7 | 13.5 | 4.5 | 8 | 38 |
| **GEM-RC** | 11.2 | 1.2 | 5.9 | 1.1 | 9 | 15 | 4.6 | 9.5 | 38 |
| **Soap20RC** | 10.5 | 1 | 5.3 | 0.7 | 8.9 | 12.7 | 4.2 | 6.8 | 44 |

Isolate TZLM3-RCL produces near-spherical cysts 8.5 – 10.5 μm (n=7) outer diameter, consisting of an internal wall 5.5-7 μm in diameter (n=7) around the eccentric cell body and an external wall separated by a space up to 2 μm, with a single conspicuous, slightly protruding plug crossing both walls (Fig 1L-M, 2T). Occasionally small particles (some resembling bacteria) can be found in the space between the two walls (not shown). Both walls remain upon excystment (Fig 1M).

There is substantial variation in size and shape within each of the four isolates within (Fig 2O) and between imaging sessions, likely depending on a combination of factors including cell density, feeding state, and age of culture. Likely cell division stages were observed, including cells with two sets of emergent flagella (Fig 2R).

### Ribosomal rRNA gene sequences and phylogenetics

SSU rDNA sequences obtained via PCR amplification and Sanger sequencing were 1642 bp, 1721 bp, and 1303 bp long for TZLM1-RC, GEM-RC, and Soap20A-RC, respectively. The genomic sequence obtained from the TZLM3-RCL assembly was 1767 bp. The SSU rDNA sequences had a GC content in the 42-46% range. No ribososomal introns were detected based on alignment and sequence lengths. The most divergent SSU rDNA sequences (of TZLM3-RCL and GEM-RC) were 88% identical to each other, markedly higher than between them and any barthelonids (Fig S3).

In the SSU rRNA gene phylogeny, metamonads are not monophyletic but appear as several moderately well supported clades, including fornicates *sensu stricto*, parabasalids, preaxostylans, *Anaeramoeba*, and barthelonids (Fig 3; Fig S1). The *Skoliomonas* isolates themselves form a fully-supported clade, branching as sister to Barthelonids with 61% bootstrap support (Fig 3). *Skoliomonas* sp. isolate TZLM3-RCL forms the deepest branch, with GEM-RC as sister to TZLM1-RC and Soap20-RC. We informally name the clade containing *Skoliomonas* spp. as ‘skoliomonads’. The concatenated SSU-LSU rRNA gene phylogeny infers a clade containing fornicates, barthelonids, and skoliomonads with 76% non-parametric bootstrap support, the latter two in turn forming a clade with 60% bootstrap support (Fig S2). The ribosomal rRNA gene trees are consistent with phylogenomic results in (Williams et al., 2023), which show full support for the barthelonid+skoliomonad clade.

**Figure 3.**
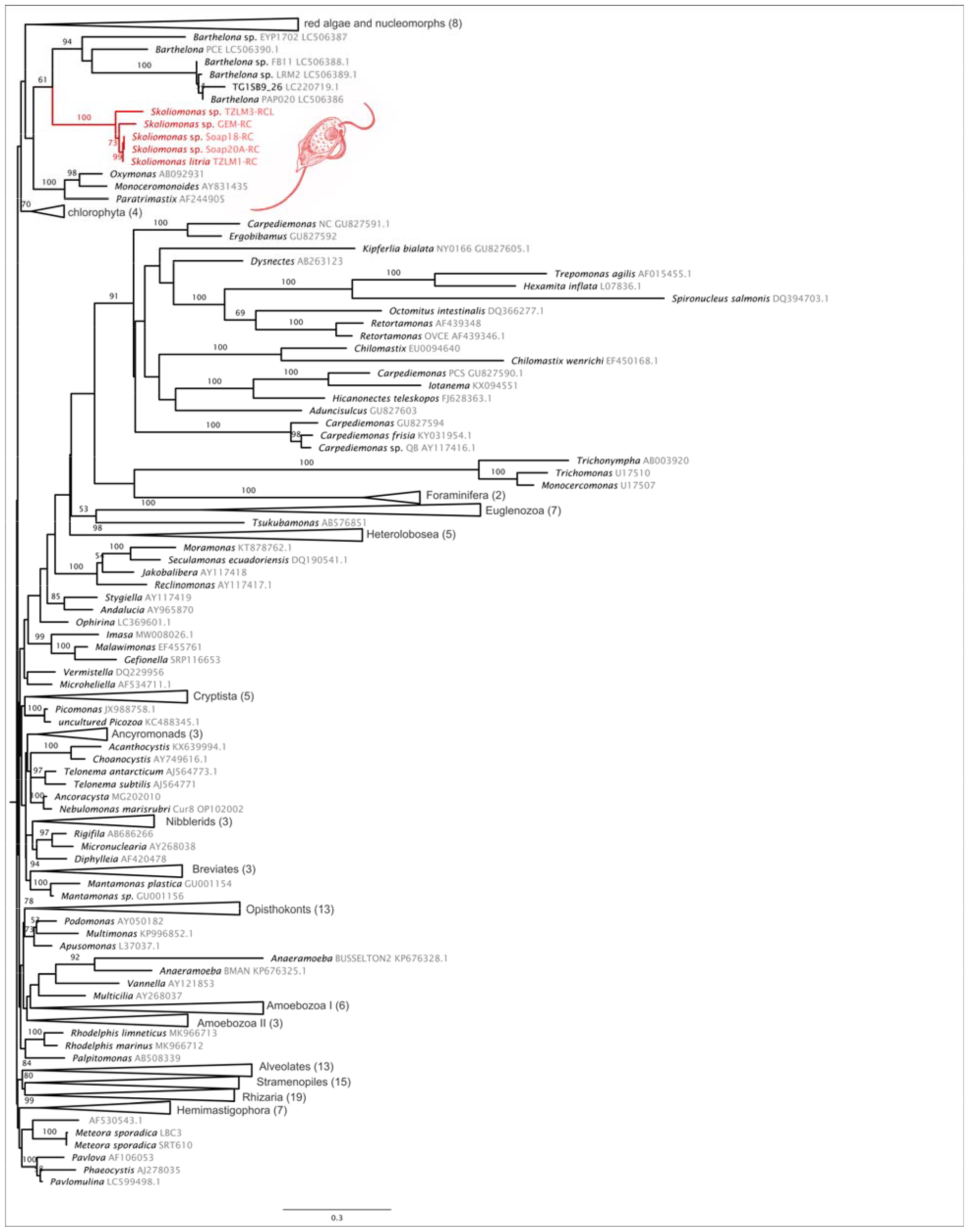
Eukaryote-wide SSU rRNA gene phylogeny inferred under the GTR+Г model in RAxML. Number in brackets indicate number of taxa in collapsed clades. Numbers on branches represent non-parametric bootstrap support values (>50%). For the complete figure with expanded collapsed clades see Figure S1.

**Figure 4:**
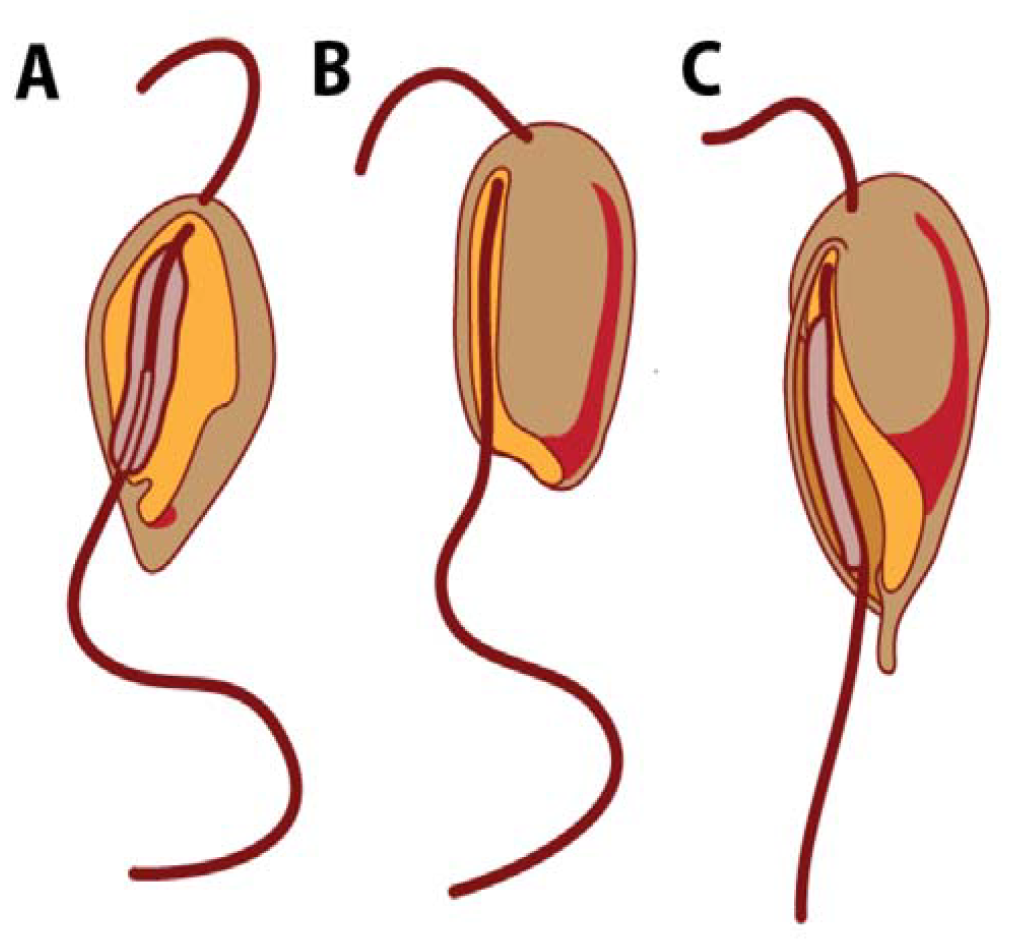
Comparison of the groove morphology between CLOs (*Carpediemonas*) (A), barthelonids (B), and skoliomonads (C). Ventral groove is indicated in orange, the cytopharynx in red, and flagellar vanes in puce.

## Discussion

The four flagellate cultures we established from three geographically disparate alkaline lakes represent a novel metamonad lineage. Phylogenetic analyses place this lineage as sister to *Barthelona* spp. (‘barthelonids’), a group of anaerobic flagellates isolated predominantly from marine environments (Bernard et al., 2000; Yazaki et al., 2020). Barthelonids have a long posterior flagellum lacking a vane discernible under light microscopy, a ventral groove in the form of a narrow channel that twists slightly left towards the posterior, and, most notably, a conspicuous posterior feature formed by the opening of the cytopharynx that curves back up along the dorsal side of the cell (Bernard et al., 2000; unpublished data). The ventral groove in *Skoliomonas* spp. is somewhat wider and less channel-like, but the open posterior as well as conspicuous cytopharyngeal element are shared in the two lineages. The right wall of the groove is much broader in skoliomonads than in barthelonids, where are distinct ‘lip’ is not observed. In contrast, most *Carpediemonas*-Like Organisms (CLOs) have a wide ventral groove (Park et al., 2010; Simpson and Patterson, 1999; Yubuki et al., 2016, 2013), as do trimastigids (O’Kelly et al., 1999; Simpson et al., 2000; Zhang et al., 2015).

### Cytopharynx

In free-living ‘typical excavates’, the posterior end of the groove hosts the site of particle ingestion featuring the opening of the cytopharynx. The cytopharynx is barely separate from the groove in some taxa (eg. *Carpediemonas (Simpson and Patterson*, 1999)), but in others extends a short distance, such as in the CLOs *Hicanonectes* (Park et al., 2009a), *Aduncisulcus* (Yubuki et al., 2016), and *Kipferlia* (Yubuki et al., 2013), as well as the trimastigids *Trimastix* (Zhang et al., 2015) and *Paratrimastix* (Simpson et al., 2000). The J-shaped cytoskeletal structure observed in Barthelonids (Bernard et al., 2000; Yazaki et al., 2020) may represent a similar cytopharynx, but considerably longer and extending back around the dorsal side of the cell almost to the flagellar insertion site. In *Skoliomonas* isolates, the cytopharynx likewise extends along the entire dorsal side of the cell, curving gently to the left. The extent to which the cytopharynx extends back up the dorsal side of the cell in barthelonids and skoliomonads is unusual among metamonads. The most similar case is in the predominantly commensal retortamonads, which have an elongate cytopharynx at the posterior of the groove that curves leftwards and then anteriorly (Bernard et al., 1997; Kulda et al., 2017); this extension is most notable in free-living *Chilomastix cuspidata*, though the cytopharynx does not extend all the way up the anterior (Bernard et al., 1997; Weerakoon et al., 1999).

The large opening to the cytopharynx, taking up nearly half of the groove, is distinctive of skoliomonads. Altogether, the short and discreet left-pointing posterior cytopharynx in CLOs and its substantial counterpart in skoliomonads and barthelonids imply the presence of a cytopharynx in the fornicate last common ancestor, although whether the reduction in the former or the elongation in the latter are derived remains unresolved.

### Flagellar vanes

Flagellar vanes, in particular a broad ventrally-facing vane, are commonly distributed across metamonads; for example, CLOs *Aduncisulcus* (Yubuki et al., 2016) and *Hicanonectes* (Park et al., 2009a) have one prominent ventral vane (the latter has a subtle dorsal vane only detectable in TEM); *Dysnectes* (Yubuki et al., 2007) and *Kipferlia* (Yubuki et al., 2013) have both dorsal and ventral vanes, while *Carpediemonas* (Simpson and Patterson, 1999) has also an additional short lateral vane. Trimastigids likewise have both ventral and dorsal vanes (Brugerolle and Patterson, 1997; O’Kelly et al., 1999; Simpson et al., 2000; Zhang et al., 2015). Barthelonids do not appear to have flagellar vanes visible by light microscopy (Bernard et al., 2000; Yazaki et al., 2020; unpublished data), but there are no published TEM studies on the flagellar structure of the group. The presence of a broad, conspicuous ventral-facing vane in skoliomonads is consistent with the vanes in CLOs and other metamonads, and suggests a secondary loss in barthelonids.

Recent work has shown a close relationship between the flagellar vane organisation and flagellar movement, and it is predicted, based on modelling, that the vane may improve the energy efficiency of feeding current generation when operating within the groove (Suzuki et al., 2023; Suzuki-Tellier et al., 2024). Whether there is a further relationship with the type (e.g. shape) of prey preferentially ingested remains to be investigated, and vaned skoliomonads with vane-less barthelonid relatives would be excellent candidates to examine what effect the flagellar vane may have on the ingestion process.

### Cysts

One isolate of skoliomonads, TZLM3-RCL, forms complex double-walled cysts with a conspicuous plugged pore. Complex cysts with pores are distinctive of some lineages of tetramitid heteroloboseans (Pánek and Čepička, 2012; Park et al., 2009b), but that group is only distantly related to skoliomonads. Most well-known cysts among metamonads are perhaps those of diplomonads such as *Giardia*, albeit with simple external structure and often fully-flagellated stages inside (Einarsson and Svärd, 2015; Erlandsen et al., 1989; Feely, 1988). Similar cysts are also known among gut-endobiotic retortamonads (Boeck, 1921; Kulda et al., 2017). In cultures of CLOs, cysts have been reported relatively infrequently; notable exceptions include *Hicanonectes* (Park et al., 2009a) and *Iotanema* (Yubuki et al., 2017). However, unlike in skoliomonads, the above-mentioned cysts are not double-walled nor do they contain pores with plugs. Cysts and pseudocysts with similarly simple external morphology are also known in free-living and parasitic trichomonads (Céza et al., 2022; Farmer, 1993; Pereira-Neves et al., 2003), and have been observed in one isolate of trimastigids (O’Kelly et al., 1999). To our knowledge, skoliomonad pored cysts are unique among metamonads, but data on cyst morphology in this group—particularly among free-living lineages—are currently patchy and likely incomplete.

### Further directions

Ultrastructural studies of *Skoliomonas* spp., coupled with those of barthelonids, would be needed to better characterise ‘excavate’ groove morphology and evolution, particularly of the last metamonad common ancestor. Given the position of the skoliomonad+barthelonid clade as sister to fornicates (Williams et al., 2023), ultrastructural studies would help characterise the nature of the fornicate last common ancestor as well. Feeding, from prey selection to the role of the flagellar vane in particle ingestion, would be an additional topic to study in relation to groove ultastructure.

Barthelonids are anaerobes with substantial mitochondrial reduction (Yazaki et al., 2020); as a representative of a major metamonad lineage, *Skoliomonas* spp. presents an opportunity for further study of adaptation to anoxia among metamonads, e.g. by searching for anaerobic metabolism and MRO associated genes in their genomes (Williams et al., 2023). Coupled with genomics, the established cultures of *Skoliomonas* spp. could provide an excellent model system for exploring polyextremophily in eukaryotes: searching for molecular signatures of adaptations to high salinity and high pH environments, unusually within the background of adaptation to anaerobiosis. Furthermore, the eukaryotes in these cultures doubtlessly interact directly and indirectly with the prokaryotic community around them. It would be interesting to investigate the biochemical underpinnings of these relationships and compare them with those of mesophilic systems.

### Taxonomic Summary

*Skoliomonas* Eglit & Simpson n. gen.

Description: Biflagellate hunchbacked cell with a posterior spike up to half a cell body length long. A ventral groove extends the length of the cell and curves to the left towards the posterior. A wide, thin sheet forms the right wall of the groove. The distal half of the groove hosts the opening to a conspicuous recurrent cytopharynx, which tapers as it extends back up along the dorsal end of the cell. Bacterivorous anaerobe. The posterior flagellum bears a vane facing away from the groove, wide enough to be discernible by light microscopy. Some isolates form a double-walled cyst traversed by a single conspicuous plug.

### Etymology

From σκoλιóς (skolios) = bent, crooked, referring to the hunched over appearance of the cell and the twisted ventral groove, and μoνάς (monas) = unit, common suffix used for single-celled protists.

### Type species

*Skoliomonas litria* (see below)

### Zoobank registration

Described under the Zoological Code; Zoobank registration (to be added upon publication)

*Skoliomonas litria* Eglit & Simpson n. sp.

### Description

Biflagellate hunchbacked cell 8-15 μm long and 6-9 μm wide, with a posterior spike up to half a cell body length long. A ventral groove extends the length of the cell and curves to the left towards the posterior. A wide, thin sheet forms the right wall of the groove. The distal half of the groove hosts the opening to a conspicuous cytopharynx, which tapers as it extends back up along the dorsal end of the cell. Numerous sizeable food vacuoles appear in the left anterior half of the cell, filled with bacteria. The posterior flagellum bears a vane facing away from the groove, wide enough to be readily discernible by light microscopy. Haloalkaliphilic anaerobe.

### Type material

The name-bearing type (hapantotype) is an osmium-fixed resin embedded permanent slide of isolate TZLM1-RC, deposited at (to be added upon publication) (see Fig S4). This material also contains uncharacterised prokaryotes, explicitly excluded from the hapantotype.

### Type locality

Lake Manyara, Tanzania (3°37’01.7”S, 35°44’22.8”E), 16% salinity pH ∼10

### Etymology

From Ancient Greek λLτρoν (‘litron’), an alternative form of νίτρoν (‘nitron’, sodium carbonate) thought to be borrowed from Ancient Egyptian, used by Herodotus (Hdt. 2.86,87) to describe embalming salts used for mummification in Ancient Egypt, which were harvested from the carbonate-rich soda lakes of the Natron Valley (Wadi El-Natrun), Egypt. We chose the African-specific version of the Ancient Greek word to reflect the type locality and to consider possible relevance of alkaline lakes to the human culture and history around them.

λίτρoν (litr(on)) + -ια (ia) (feminine adjective-forming suffix, “of”)

### Gene sequence

The SSU rRNA gene from isolate TZLM1-RC has the GenBank accession number PP416847

### Zoobank registration

Described under the Zoological Code; Zoobank registration TBA

## Acknowledgements

We’d like to thank Z. Nathan Taylor for obtaining the samples for isolates TZLM1-RC and TZLM3-RCL, as well as Jackie Zorz (University of Calgary) and Ashley R. Smith for samples for GEM-RC and Soap20-RC, respectively. We’d also like to thank Aaron Beek (North-West University and Case Western Reserve University) for consultation on Ancient Greek for the taxonomic names. This work was supported by an NSERC Discovery grant 298366-2019 awarded to AGBS.

## Data availability

SSU rRNA gene sequences have been submitted to GenBank under accession numbers PP416847 - PP416851. Cultured isolates available upon request.

