## Supplementary figures for "Characterisation of *Skoliomonas* gen. nov., a haloalkaliphilic anaerobe related to barthelonids (Metamonada)"

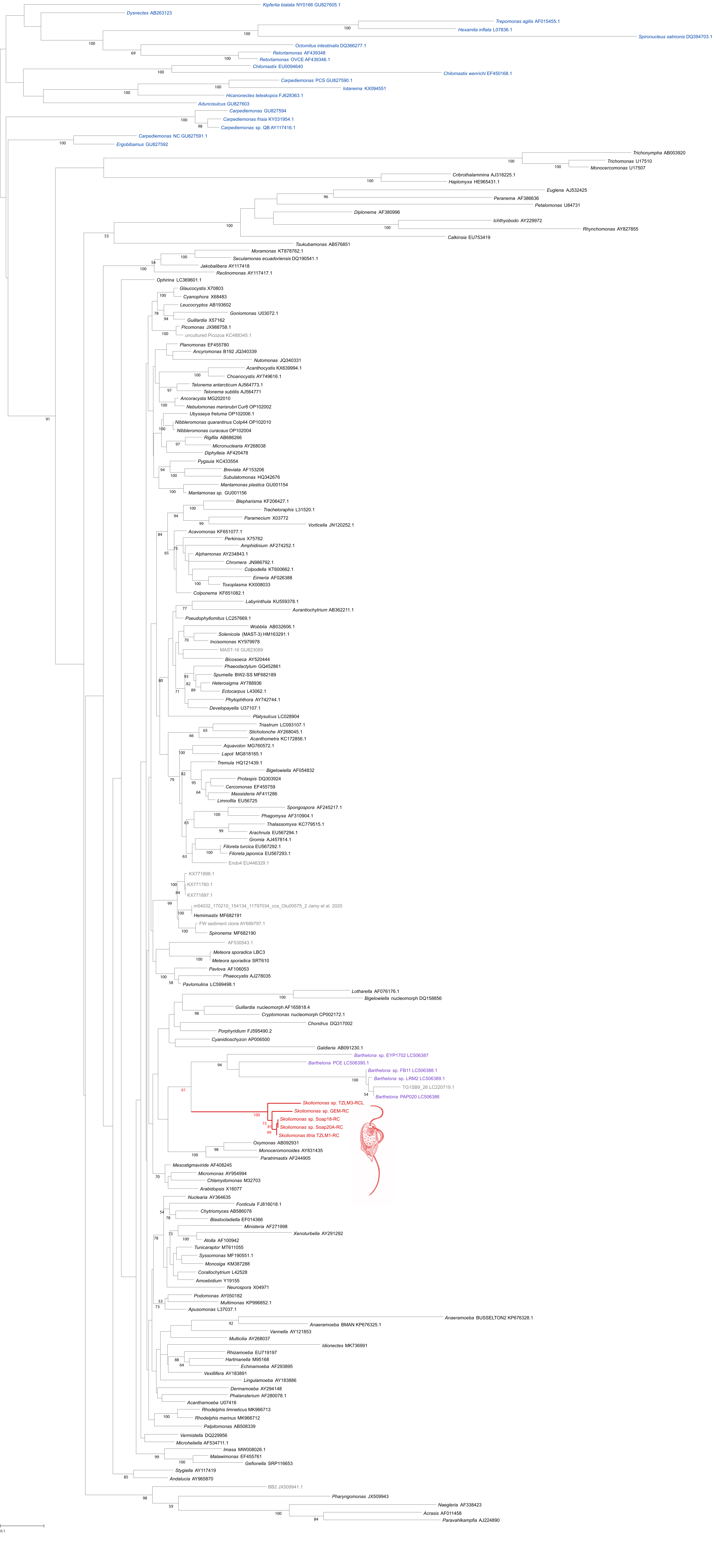

**Figure S1.** Eukaryote-wide SSU rRNA gene phylogeny (1321 sites x 219 taxa) inferred under the GTR+Г model in RAxML. *Skoliomonas* spp. isolates highlighted in red, Barthelona spp. in violet, and fornicates are indicated in blue. Support values under 50% (non-parametric boostrap, 500 replicates) omitted for clarity.

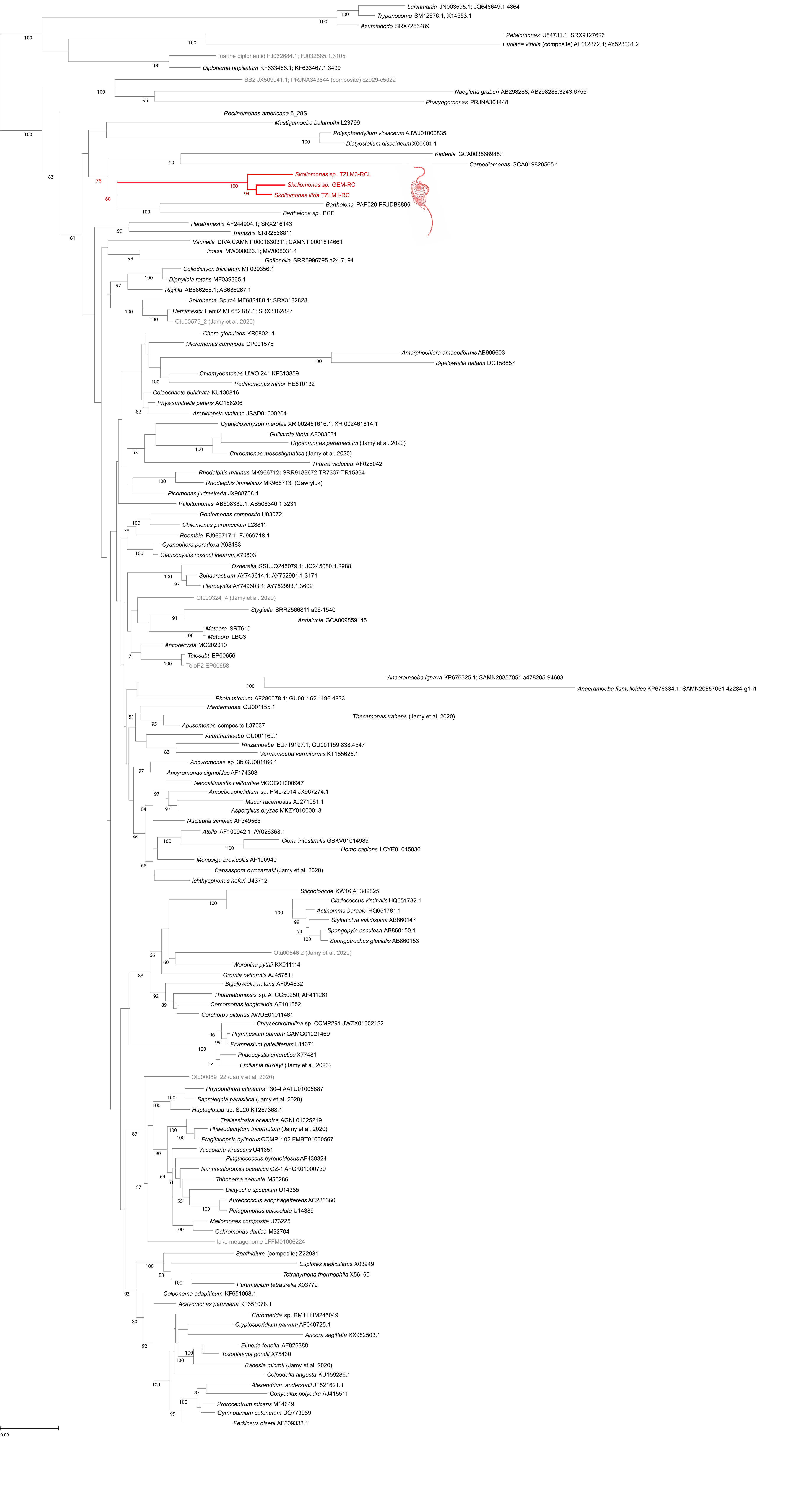
**Figure S2**. Concatenated SSU-LSU rRNA gene phylogeny representing eukaryote-wide diversity inferred under the GTR+Г model in RAxML from 3064 sites and 141 taxa. *Skoliomonas* spp. isolates highlighted in red. Support values under 50% (non-parametric bootstrap, 500 replicates) omitted for clarity.

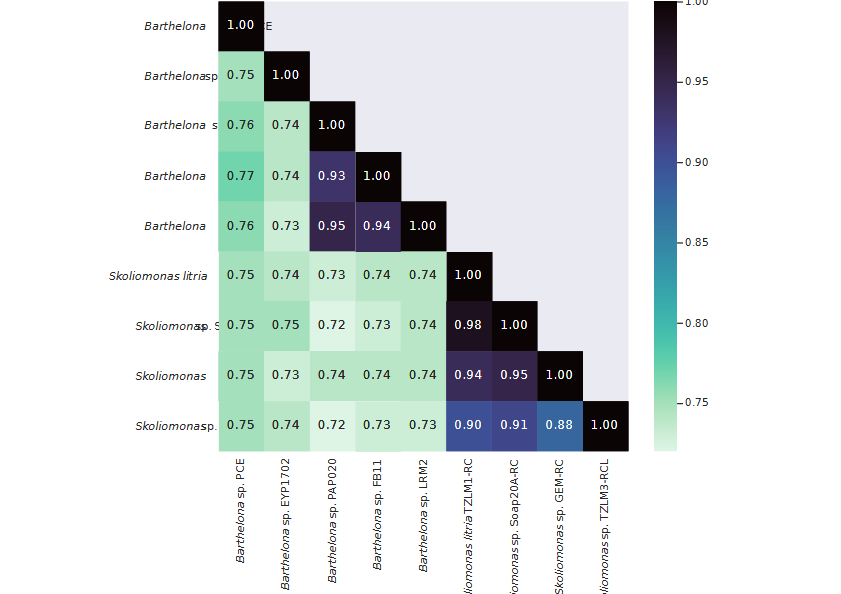

**Figure S3**. Distance matrix of sequence identities between SSU rDNA sequences of the skoliomonad and barthelonid isolates.

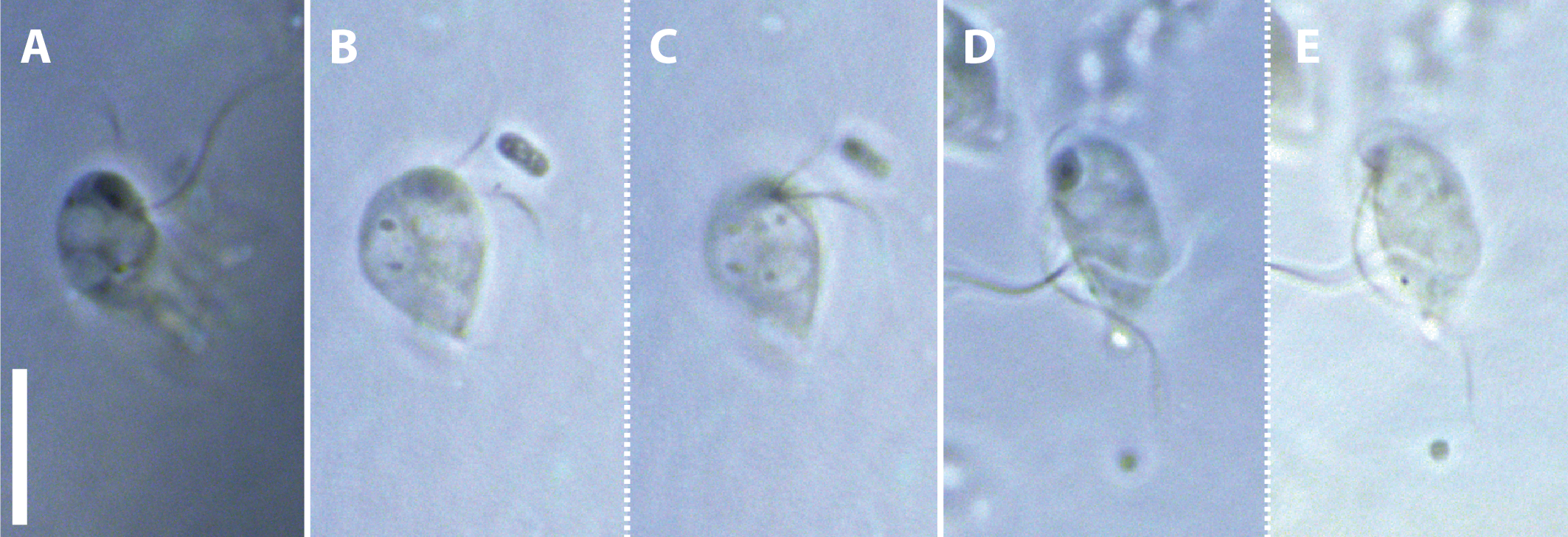
**Figure S4**. Phase contrast light micrographs of representative cells in the hapantotype preparation of resin-embedded osmium-fixed cells on a slide. B,C and D,E are each a pair of optical sections of the same cell. Note the dark-staining nucleus and nucleolus in A, B and D, and the conspicuous right margin of the groove in C and E. Scale bar = 10 µm.
